# TCGA Expression Explorer: A three-generation comparison of TCGA RNA-seq reveals pipeline-dependent loss of genes without unique exonic sequence

**DOI:** 10.64898/2026.11.25.769128

**Authors:** Shoichiro Tange, Masashi Idogawa

## Abstract

**Background:** Expression values for The Cancer Genome Atlas (TCGA) have been updated twice, so three generations now exist: the legacy release (RSEM, hg19), Genomic Data Commons harmonization (GDC v1; HTSeq-count, GENCODE v22), and the 2022 re-processing (GDC v2; STAR GeneCounts, v36). All three remain in use, differ in identifiers, units, and gene content, and no interface shows one gene across them.

**Findings:** TCGA Expression Explorer is a static browser tool that draws one gene’s distribution in five cancer-type cohorts under all three generations. We compared detection across specimens and the 17,352 genes common to them, measuring the fraction of each gene’s exonic bases that also lie in another gene’s exons in the annotation of that generation. Of the genes detected in the legacy release, those becoming zero in every GDC v2 specimen were rare below 90% overlap (at most 0.5%), occasional at 90%–<100% (0%–21%), and universal at 100%: 46–55 genes per cohort with no exonic sequence of their own. The same class is 82%–85% in GDC v1 owing to its counting rule. Seventy genes are undetectable in both GDC generations and 32 in all five cohorts, among them *TREX1*, whose exons are entirely covered by *ATRIP* in v36 and by an *ATRIP*–*TREX1* readthrough in v22. Counting public alignments in six ways reproduces the loss.

**Conclusions:** For a predictable set of genes, measurability depends on the processing generation. Discarding reads compatible with more than one gene suffices to cause the loss, and the affected genes can be predicted from the annotation alone: https://5spgm.github.io/tcga-expression-explorer/.

## Background

Two decades after its launch, The Cancer Genome Atlas (TCGA) remains one of the most heavily used resources in cancer research [1]. However, its RNA-seq data are not a single dataset but three, produced by three pipelines between 2010 and 2022. All three are still downloaded, cited, and reused. We have worked with these data ourselves (in studies of a candidate oncogene in pancreatic cancer [2] and of a long non-coding RNA modulating p53 function in colorectal cancer [3]). We have argued before that TCGA analysis is only as sound as the user’s attention to how the underlying dataset is assembled, pointing out a pitfall of survival analysis with TCGA data that is invisible unless the clinical annotation is examined directly [4]. The present study applies the same attention to the expression matrices.

The legacy release, distributed by the TCGA DCC until the data migrated to the NCI Genomic Data Commons [5], was produced by the UNC RNASeqV2 pipeline: MapSplice alignment to hg19 [6] followed by RSEM quantification [7] against a UCSC-derived gene model of 20,531 genes, reported as `normalized_count`, an upper-quartile-scaled RSEM estimate rather than a raw read count. The first GDC harmonization realigned the raw reads to GRCh38 with STAR [8] and counted them using HTSeq count [9] against GENCODE v22, yielding 60,483 genes as counts, FPKM, and FPKM-UQ. In 2022, the GDC replaced that workflow with STAR’s own gene counter (`--quantMode GeneCounts`) against GENCODE v36, yielding 60,660 genes and adding TPM. Table 1 summarizes these three parameters.

**Table 1.** Three generations of TCGA gene-level RNA-seq.

|  | **Legacy DCC** | **GDC v1** | **GDC v2 (STAR-Counts era)** |
|---|---|---|---|
| Period | 2010–2016 | 2016–2022 | 2022– |
| GDC data releases | n/a (TCGA DCC Data Matrix) | harmonized releases up to DR-31.0 | DR-32.0 onwards |
| Reference genome | hg19 | GRCh38 | GRCh38 |
| Alignment | MapSplice | STAR 2-pass | STAR 2-pass |
| Quantification | RSEM | HTSeq-count, intersection-nonempty, unstranded | STAR GeneCounts, unstranded |
| Gene annotation | UCSC hg19 gene model | GENCODE v22 | GENCODE v36 |
| Genes in matrix | 20,531 | 60,483 | 60,660 |
| Unit used here | normalized count | FPKM-UQ (FPKM for BRCA) | TPM |

None of these releases superseded the others in practice. The legacy matrices persist in mirrors and in the supplementary tables of a decade of publications, the GDC v1 matrices persist in widely used access layers and compendia [10,11], and independent reprocessing of the same reads also exists [12,13,14]. A figure legend that reads “expression of gene *X* in TCGA” therefore does not tell the reader which generation produced the value, or a reader who wants to check faces three matrices that differ in identifier system (gene symbol with Entrez ID, versioned Ensembl gene ID, Ensembl gene ID with symbol), in unit, and in which genes they are present at all.

The legacy data have been compared with the harmonized data before but not with GDC v2. Gao et al. [15] performed a systematic legacy-versus-harmonized comparison across five TCGA data platforms and concluded that the hg19 and hg38 datasets are highly concordant; for mRNA, they used 2,302 samples from three cohorts and the 19,744 protein-coding genes that mapped unambiguously between the two releases and found the per-sample rank correlation of expression well preserved (mean Spearman’s rho 0.943, range 0.893–0.959). The GDC’s own description of harmonized workflows [16] and studies on how the choice of gene annotation affects RNA-seq quantification [17] also report a close agreement between generations. Our results are consistent with these findings. The rank correlation reported by Gao et al. [15] measures the agreement over the genome as a whole, and a few dozen genes falling to zero barely move it. Their restriction to genes that are unambiguously mapped between releases also removes a part of the overlapping class studied here before any values are compared. Moreover, their comparison predates the 2022 re-processing by three years and therefore ends at GDC v1, and the authors themselves call for “similar comparisons as new data emerge and reference data evolve” [15]. To our knowledge, no published comparison has yet included GDC v2; the present study provides one such comparison.

The pipeline and annotation choices that change RNA-seq values are well established: (i) the SEQC/MAQC-III benchmarking established the magnitude of protocol and pipeline effects [18], (ii) multi-mapping and ambiguous reads are known to affect disease-relevant gene families [19], and (iii) transcript-level quantification followed by gene-level aggregation is known to yield different (and generally better-behaved) gene-level estimates than direct union counting [20,21]. What is missing in day-to-day practice is not the knowledge but the check: nothing in the usual workflow tells a user that the particular gene in front of them is one where the generations disagree.

We built TCGA Expression Explorer to make that check trivial, and we report here the systematic effect that the tool surfaced first: for genes whose exons are entirely claimed by another gene, the re-processed generations do not merely shift the values; they remove the gene from the measurable set.

### Data collection, curation, and quality control

The labels “GDC v1” and “GDC v2” are our shorthand; in the GDC’s own terms, they are the HTSeq-era releases up to Data Release 31.0 and the STAR-Counts-era releases from Data Release 32.0, the release in which the HTSeq pipeline was retired and gene-level normalized counts, FPKM, FPKM-UQ, and TPM were consolidated into the STAR-Counts files [22]. Legacy gene-level matrices (`normalized_count`, UNC Illumina HiSeq RNASeqV2) were obtained from the TCGA DCC Data Matrix bulk distribution, which predates GDC release numbering, between April 2014 and August 2015, depending on the cohort. The GDC v1 matrices (FPKM-UQ) were downloaded from the GDC Data Portal [23] on February 5, 2021, that is, Data Release 28.0, and the BRCA GDC v1 matrix (FPKM) was used instead of the UCSC Xena GDC hub [10]. GDC v2 matrices (TPM) were downloaded per cohort at different times and therefore from different releases: Data Release 36.0 for COAD and PAAD, 41.0 for STAD, 43.0 for BRCA, and 46.0 for LUAD. The exact file manifests, with download dates and file checksums, are part of the deposit [24]. Table 2 lists the release behind every matrix. A single preprocessing script (`scripts/preprocess_tcga.py` in the source repository [25]) reads the three matrices of one cohort and writes one JSON document per gene containing the tumor and normal value vectors of each generation, together with the unit label and specimen counts.

**Table 2.** Samples displayed by the tool and the provenance of each matrix.

| **Cohort** | **Legacy DCC: specimens (archive, downloaded)** | **GDC v1: specimens (release)** | **GDC v2: specimens (release)** | **Specimens compared in all three** |
|---|---|---|---|---|
| BRCA | 1,147 (1.8.0, Apr 2014) | 1,197 (UCSC Xena GDC hub, Apr 2024) | 1,205 (DR-43.0, Jun 2025) | 1,138 |
| COAD | 313 (1.8.0, Aug 2014) | 497 (DR-28.0, Feb 2021) | 499 (DR-36.0, Dec 2022) | 313 |
| LUAD | 546 (1.13.0, Sep 2014) | 572 (DR-28.0, Feb 2021) | 576 (DR-46.0, Aug 2026) | 543 |
| PAAD | 34 (1.8.0, Jul 2015) | 153 (DR-28.0, Feb 2021) | 154 (DR-36.0, Feb 2023) | 33 |
| STAD | 450 (1.0.0, Aug 2015) | 407 (DR-28.0, Feb 2021) | 447 (DR-41.0, Oct 2024) | 403 |
| **Total** | **2,490** | **2,826** | **2,881** | **2,430** |

Tumor and normal specimens, after the exclusion of formalin-fixed and cohort-flagged specimens and collapsing of duplicate vials. Metastatic and recurrent specimens were also excluded. Legacy matrices predate GDC release numbering; hence, the download date is given, The PAAD legacy matrix contains 34 specimens because that cohort was still accruing when its legacy snapshot was taken in July 2015, which is also why the legacy row of that cohort is the smallest specimen set in the comparison; for the GDC generations the release is the one current at the download date recorded in the deposited file manifests. The BRCA GDC v1 matrix came from the UCSC Xena GDC hub rather than the portal.

Specimen selection is identical for all generations of a cohort: barcodes are reduced to TCGA sample level, and primary tumor and solid normal specimens are retained; metastatic and recurrent specimens are excluded by default and can be folded in from the interface; duplicate vials of one patient are reduced to the first vial; and formalin-fixed specimens and cohort-specific flagged specimens are removed using exclusion lists generated by `scripts/make_ffpe_exclude_list.py` and deposited with the data. Subtype assignments were converted to a common two-column format using `make_*_subtype_table.py` scripts from the supplementary tables of the respective publications [26,27,28,29] and the Biospecimen Core Resource clinical patient file for triple-negative status.

### User interface

TCGA Expression Explorer [30] (Fig. 1) is a static web application. The user selects a cohort, types a gene symbol or Ensembl identifier, and receives three panels, one for each generation (legacy DCC, GDC v1, and GDC v2). Each panel shows the distribution of the gene’s values in the tumor and matched normal specimens of the same cohort, with the unit of that generation on the axis and the number of specimens in each box printed beneath it. Tumor specimens can additionally be split by any of 11 groupings (nine published molecular classifications together with immunohistochemistry-based triple-negative status and a tumor-purity class), of which two to six apply to a given cohort, so that a generation difference can be distinguished from a subtype difference: pan-cancer iCluster [26]; molecular subtype, microsatellite-instability status, CpG-island-methylator phenotype, and colorectal consensus molecular subtype [27]; immune subtype [28]; the Moffitt and Bailey pancreatic subtypes and tumor purity class [29]; PAM50 as assigned by the TCGA breast cohort; and triple-negative status derived from the Biospecimen Core Resource clinical files. A log₂(*x* + 1) toggle; an option to fold metastatic and recurrent specimens back in; and PNG, SVG, and CSV export of the displayed data are provided.

**Figure 1.**
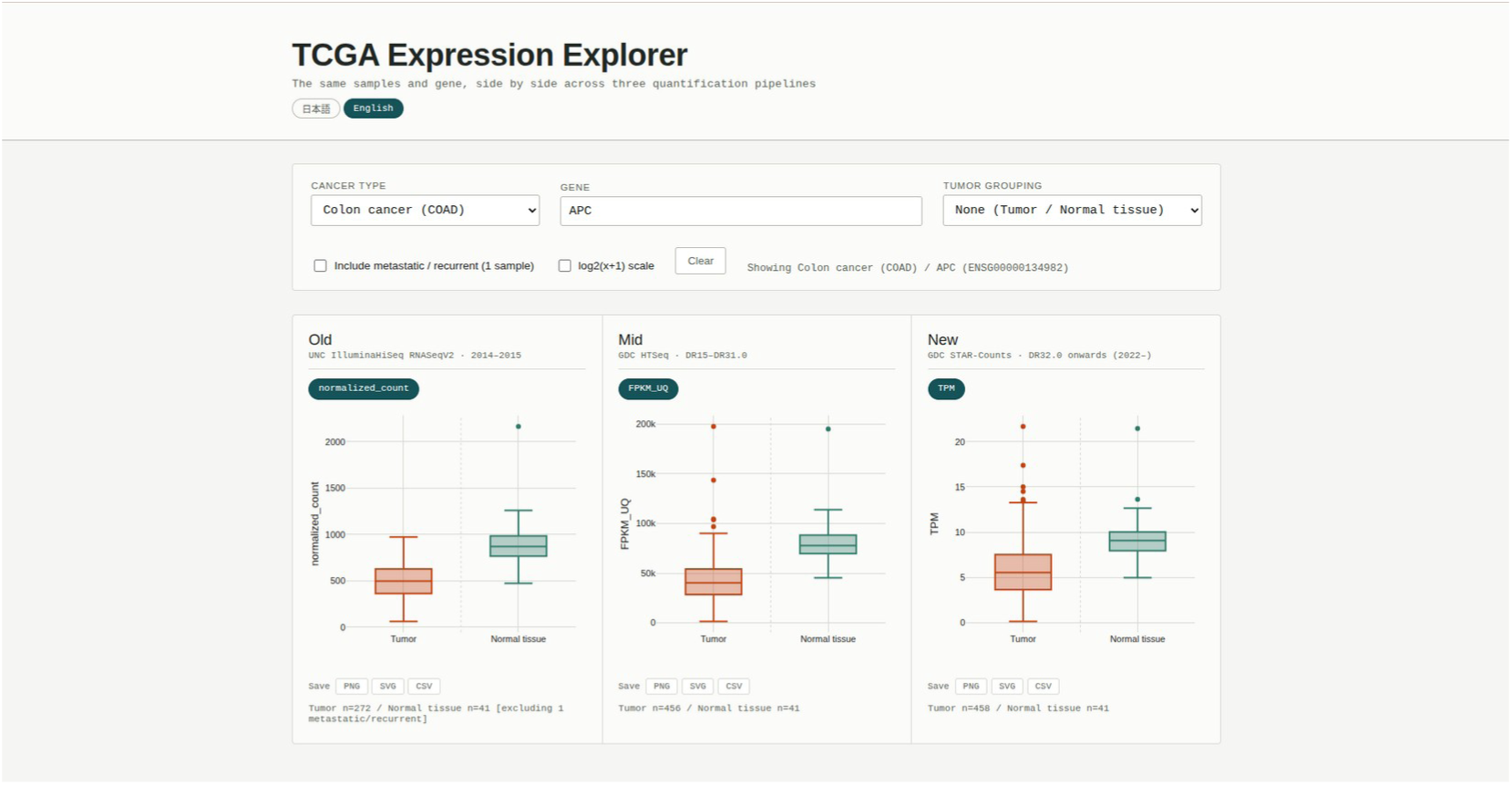
TCGA Expression Explorer interface. A single query (cohort, gene, optional tumor grouping) returns the distribution of that gene in the same cohort under all three quantification generations: legacy DCC (UNC IlluminaHiSeq RNASeqV2, `normalized_count`), GDC v1 (HTSeq era, FPKM-UQ), and GDC v2 (STAR-Counts era, TPM). The three panels are headed *Old*, *Mid,* and *New* in the interface and carry each generation’s own source and release beneath the heading. Here, *APC* is shown for COAD with no grouping selected; thus, the boxes hold the full comparable sets of Table 2 (313, 497, and 499 specimens); tumors can instead be split by any of the available subtype schemes, in which case, only tumors carrying an assignment in that scheme are drawn. Values are shown on their native scales, each generation in its own unit; a log_2_(*x* + 1) option is available. Per-box specimen counts are printed beneath each panel, and the displayed data can be exported as PNG, SVG, or CSV.

The application performs no computation on the server. The front end is plain JavaScript, with Plotly for rendering, and the data are precomputed as one JSON document per gene per cohort (60,660 genes × 5 cohorts = 303,300 documents; the gene set is that of GENCODE v36, with the legacy and GDC v1 values mapped onto it) served as static files from object storage. There is no login, installation, or query quota. The interface is available in English and Japanese.

### Implementation

The front end is dependency-light JavaScript (no framework) with Plotly.js 2.35.2 for rendering, served from GitHub Pages. The per-gene JSON documents and a manifest are served as static files from Cloudflare R2 object storage, and the data location is a single configurable constant. Gene lookup uses a per-cohort index file; therefore, a query fetches one index and one gene document.

### Intended uses and benefits

This tool is aimed at three situations that arise routinely. A reader who meets a TCGA expression claim in a paper can see, in one screen, whether that claim survives re-quantification. An author preparing an analysis of a particular gene can check whether the choice matters for that gene before committing it to a generation. A reviewer can verify a submitted figure against the other two generations without downloading anything. In each case, the benefit is that the comparison is available at the moment the question arises, rather than requiring three downloads with three identifier systems. Because tumor specimens can be split by published subtype schemes, a difference between generations can be distinguished from a difference between subtypes.

We built on the data and annotation produced by others, and the TCGA Research Network generated the sequencing data and legacy quantification. The Genomic Data Commons produced both harmonized re-processings [5], the UCSC Xena team distributes the GDC v1 matrices in a form that can be downloaded per cohort [10], GENCODE provides the annotation on which the later generations depend [31], and the subtype assignments come from the publications cited above [26,27,28,29]. The contributions of this work are the aligned three-generation view, the curation that makes the three comparable, and the analysis in *Case study*, which identifies the genes whose measurability depends on the generation, as listed in Supplementary Table S1.

### Comparison with functionally similar resources

To establish what already exists, we searched OpenAlex for web resources that display TCGA gene-level expression and for studies comparing TCGA processing generations (queries and dates are given under *Identification of comparable resources*). The resources screened are listed in Supplementary Table S2, and the functional comparison is shown in Table 3.

**Table 3.** Interfaces displaying TCGA gene-level expression.

| Resource | TCGA generation served | Generations per gene in one view | Per-gene distribution plot | Tumor stratification | Coding required |
|---|---|---|---|---|---|
| GDC Data Portal | GDC v2 (STAR-Counts) | 1 | Yes | No | No |
| cBioPortal | Study-dependent; TCGA sets are Firehose Legacy and PanCancer Atlas | 1 | Yes | By clinical attribute | No |
| TCGEx | Inherited from cBioPortal | 1 | Yes | By clinical attribute | No |
| GEPIA2 | None of the three; UCSC Xena Toil recompute | 1 | Yes | By stage, tumor vs normal | No |
| UALCAN | GDC v1 (HTSeq-FPKM workflow) | 1 | Yes | By clinical subgroup | No |
| OncoDB | None of the three; own STAR + RefSeq pipeline | 1 | Yes | By clinical variable | No |
| TNMplot | GDC v1 (HTSeq-Counts, DESeq2-normalized) | 1 | Yes | Tumor/normal/metastatic | No |
| UCSC Xena | Legacy, GDC v1 and GDC v2, as separate hubs | Multiple, not aligned | Yes | By phenotype column | No |
| TCGAbiolinks, recount3 (R/Bioconductor), Toil recompute (compendium) | Any release the user downloads, or own re-processing | User-defined | User-defined | User-defined | Yes |
| **TCGA Expression Explorer** | **Legacy + GDC v1 + GDC v2, same specimens** | **3** | **Yes** | **11 groupings (9 published)** | **No** |

“Generations” refers to the quantification generations of Table 1 that a user can see for one gene without leaving the interface.

Several widely used resources answer the question “what are the expression levels of gene *X* in cancer *Y*,” and each answers it from one generation. We read the data provenance statement for each resource to establish which generation it serves (Table 3). The GDC Data Portal [23] serves GDC v2. UALCAN [32] states that its expression files were obtained from the GDC through the HTSeq-FPKM workflow and TNMplot [33], which normalizes HTSeq-Counts files with DESeq2; therefore, both serve as the GDC v1 generation. The GEPIA series [34,35,36] serves Xena-hosted data rather than any GDC release; GEPIA2 uses the UCSC Xena Toil recompute [13], the current GEPIA3 lists its expression source as the Xena TCGA PANCAN dataset version-dated 2016-09-01, and OncoDB [37] realigns the raw reads itself and summarizes them against the RefSeq transcriptome. cBioPortal [38] reveals one processed release per study; its TCGA sets were the Firehose Legacy and PanCancer Atlas matrices, and TCGEx [39] inherited these sets. GENT2 [40] serves as a microarray rather than RNA-seq data. UCSC Xena [10] is the closest, hosting both a legacy hub and GDC hubs, and its visual spreadsheet can place columns from different datasets next to each other; however, the generations are separate datasets that the user must identify by name, with different identifier systems and units and no guarantee that the columns refer to the same specimens. Programmatic access provides full flexibility at the cost of writing code for each comparison. TCGAbiolinks [11] and recount3 [14] are R/Bioconductor packages, and Toil recompute [13] is a downloadable, uniformly re-processed compendium distributed through Xena.

None of them shows one gene under more than one quantification generation, aligned on the same specimens, in a single view, and for several of them, the generation is not stated in the interface at all; it has to be recovered from the methods section of the paper, as in Table 3, or inferred from the data. This is the gap that the tool fills, and the cost of filling it is its scope: five cohorts and gene-level expression only. Thus, the tool complements rather than replaces general-purpose portals.

### Data content

Table 2 shows the number of specimens displayed per cohort and generation. Specimen sets differ between generations because the underlying releases differ; for example, PAAD is represented by 34 specimens in the legacy release and 153–154 in the GDC releases. Formalin-fixed specimens and cohort-specific flagged specimens are excluded, duplicate vials from the same patient are collapsed, and metastatic and recurrent specimens are excluded by default, as described above. The source matrices, specimen annotation, exclusion lists, and provenance records for every file are deposited [24] (see Data Availability) so that the numbers shown by the tool can be reproduced independently.

### Testing and reproducibility

The source repository [25] contains preprocessing and analysis scripts, a pinned dependency list, and a test bundle. Three input matrices of one cohort are reduced to a defined gene subset, together with the expected JSON documents and the expected output tables of the comparison reported here. Running the two commands documented in the README repository on that bundle reproduces the expected outputs byte-for-byte, which is a check that a reader or reviewer can run without downloading the full dataset. The complete input matrices, from which every number in this article can be regenerated, are in the Zenodo deposit [24], where each deposited file carries an SHA-256 checksum and a provenance record naming its source, download date, and any transformation applied. The interface requires no installation; it runs in any current browser on the project home page [30].

### Case study: genes with no unique exonic sequence are unmeasurable

The three generations reported different units (RSEM-normalized counts, FPKM-UQ, and TPM); therefore, their values cannot be compared directly. Whether a gene received any reads can be determined as follows. In all three units, zero means that no read was assigned to the gene in that specimen, and any value above zero means that some were. Therefore, we asked what fraction of specimens was non-zero for every gene comparable across the three generations. The comparison uses exactly the specimens displayed by the tool. Thus, these numbers and the distributions a reader sees in the interface come from the same selection, described under *Cross-generation comparison*: 33 specimens in PAAD to 1,138 in BRCA, a 34-fold range, and 17,352 genes in every cohort.

Both GDC counting steps assign a read to a gene only when it is compatible with exactly one annotated gene. The GDC v1 workflow ran `htseq-count -m intersection-nonempty -t exon -s no -a 10`, and the GDC v2 workflow used STAR’s GeneCounts mode. The two implement that principle differently. STAR discards a read as soon as it is compatible with a second gene and does not count reads mapped to more than one locus. `The htseq-count -m intersection-nonempty` instead assigns the read to the intersection of the gene sets at the positions it covers, discarding it only when that intersection is empty or holds more than one gene; a read that lies inside gene A along its whole length and inside gene B over part of it is therefore still counted for A [8,9,41]. The widely used featureCounts behaves the same way by default [42]. Three properties of this rule shape the analysis. It operates on **exon** features, not on gene spans; therefore, the quantity that should matter is the share of a gene’s exonic bases that other genes also claim. Because GDC treats all RNA-seq reads as unstranded to harmonize specimens [41], a gene on the opposite strand of the same exonic interval competes for reads exactly as a same-strand gene does; therefore, the share must be taken over all other genes, irrespective of strand. The competing genes are those in the annotation against which the generation in question was counted, which is not the same annotation in the two GDC generations. Therefore, we computed, for every comparable gene, the fraction of its exonic bases that are exonic in at least one other gene, separately in GENCODE v36 and GENCODE v22, as described under *Cross-generation comparison*, and asked each generation’s question with its own annotation.

The answer is categorical at the top of the range. Among the genes robustly detected in the legacy release (non-zero in more than half of the compared specimens, 14,919–15,377 genes per cohort), we asked how many had no non-zero values **in any specimen** of the generation in question. In GDC v2, classified using GENCODE v36 (Fig. 2a):

- below 50% overlap (14,446–14,887 genes per cohort): 0.01%–0.02% zero everywhere
- 50%–90% overlap (385–399 genes): 0.00%–0.50%
- 90% to just below 100% (40–42 genes): 0.0%–21.4%
- complete overlap, no exonic base of the gene unique to it (46–55 genes): **100% in every cohort, without exception.**

**Figure 2.**
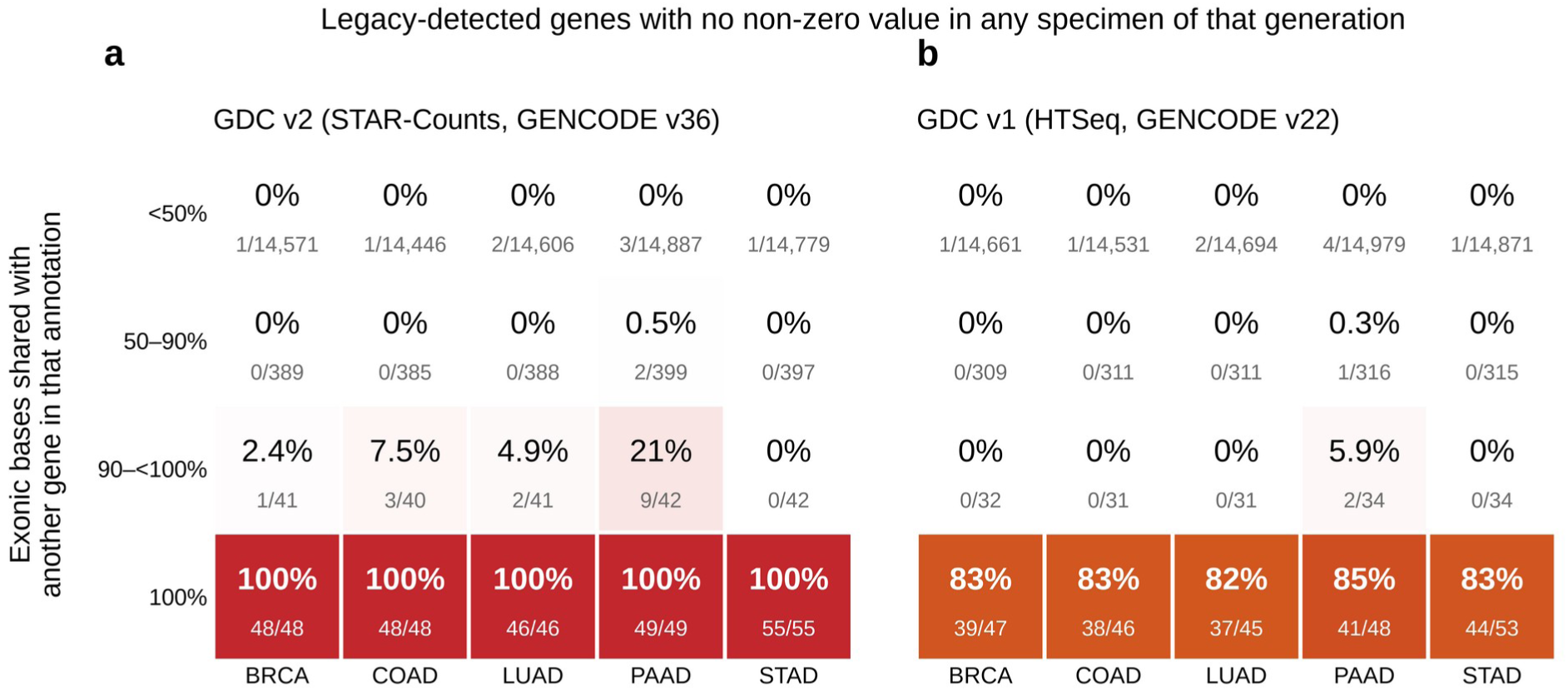
Loss of detection concentrates at complete exonic overlap in each generation’s own annotation. Each cell gives the percentage of legacy-detected genes (non-zero in more than 50% of compared specimens) that have no non-zero value in any compared specimen of that generation, with the underlying gene counts beneath. Rows are the fraction of the gene’s exonic bases that are also exonic in at least one other gene, columns are cohorts, and cells are shaded on one scale running from white at 0% to red at 100%, such that a cell at 83% appears orange. (**a**) GDC v2 (STAR-Counts), classified by GENCODE v36, the annotation it was counted against. (**b**) GDC v1, classified by GENCODE v22, its own annotation. The same counts under the weaker undetectable reading (non-zero in at most 10% of specimens) are in Table 4.

**Table 4.** Detection in each GDC generation by exonic overlap in that generation’s annotation.

| **Cohort** | **Exonic overlap in that annotation** | **GDC v2: zero everywhere** | **GDC v2: ≤10% of specimens** | **GDC v1: zero everywhere** | **GDC v1: ≤10% of specimens** |
|---|---|---|---|---|---|
| BRCA | <50% | 1/14,571 (0.0%) | 8/14,571 (0.1%) | 1/14,661 (0.0%) | 7/14,661 (0.0%) |
|  | 50%–90% | 0/389 (0.0%) | 5/389 (1.3%) | 0/309 (0.0%) | 3/309 (1.0%) |
|  | 90%–<100% | 1/41 (2.4%) | 13/41 (31.7%) | 0/32 (0.0%) | 3/32 (9.4%) |
|  | 100% | 48/48 (100.0%) | 48/48 (100.0%) | 39/47 (83.0%) | 41/47 (87.2%) |
| COAD | <50% | 1/14,446 (0.0%) | 13/14,446 (0.1%) | 1/14,531 (0.0%) | 12/14,531 (0.1%) |
|  | 50%–90% | 0/385 (0.0%) | 6/385 (1.6%) | 0/311 (0.0%) | 4/311 (1.3%) |
|  | 90%–<100% | 3/40 (7.5%) | 11/40 (27.5%) | 0/31 (0.0%) | 3/31 (9.7%) |
|  | 100% | 48/48 (100.0%) | 48/48 (100.0%) | 38/46 (82.6%) | 40/46 (87.0%) |
| LUAD | <50% | 2/14,606 (0.0%) | 9/14,606 (0.1%) | 2/14,694 (0.0%) | 7/14,694 (0.0%) |
|  | 50%–90% | 0/388 (0.0%) | 8/388 (2.1%) | 0/311 (0.0%) | 3/311 (1.0%) |
|  | 90%–<100% | 2/41 (4.9%) | 11/41 (26.8%) | 0/31 (0.0%) | 3/31 (9.7%) |
|  | 100% | 46/46 (100.0%) | 46/46 (100.0%) | 37/45 (82.2%) | 39/45 (86.7%) |
| PAAD | <50% | 3/14,887 (0.0%) | 16/14,887 (0.1%) | 4/14,979 (0.0%) | 12/14,979 (0.1%) |
|  | 50%–90% | 2/399 (0.5%) | 9/399 (2.3%) | 1/316 (0.3%) | 3/316 (0.9%) |
|  | 90–<100% | 9/42 (21.4%) | 13/42 (31.0%) | 2/34 (5.9%) | 4/34 (11.8%) |
|  | 100% | 49/49 (100.0%) | 49/49 (100.0%) | 41/48 (85.4%) | 42/48 (87.5%) |
| STAD | <50% | 1/14,779 (0.0%) | 2/14,779 (0.0%) | 1/14,871 (0.0%) | 2/14,871 (0.0%) |
|  | 50%–90% | 0/397 (0.0%) | 2/397 (0.5%) | 0/315 (0.0%) | 1/315 (0.3%) |
|  | 90%–<100% | 0/42 (0.0%) | 8/42 (19.0%) | 0/34 (0.0%) | 2/34 (5.9%) |
|  | 100% | 55/55 (100.0%) | 55/55 (100.0%) | 44/53 (83.0%) | 47/53 (88.7%) |

Rows are legacy-detected genes (non-zero in more than 50% of compared specimens) grouped by the fraction of their exonic bases that are also exonic in at least one other gene, in the annotation that generation was counted against (GENCODE v36 for GDC v2, GENCODE v22 for GDC v1). Two readings are given for each generation: genes with no non-zero value in any compared specimen, and genes non-zero in at most 10% of specimens. Figure 2 and the text use the first. The two-generation definition of *lost* used in Tables 5 and 6 is the second, applied to both GDC generations at once with the class taken from GENCODE v36.

The same question put to GDC v1 with GENCODE v22 gives the same shape with a softer top: 0.01%–0.03% below 50% overlap, 0.00%–0.32% at 50%–90%, 0.0%–5.9% at 90%–<100%, and 82.2%–85.4% of the 45–53 genes per cohort with complete overlap (Fig. 2b). Therefore, the effect is concentrated at a complete overlap rather than spread across the range. Below that threshold, the proportions are small, and the ordering they show tracks the number of specimens rather than the degree of overlap: in the 90%–<100% class of GDC v2, the strict figure is 2.4% in BRCA (1,138 specimens), 4.9% in LUAD (543), 0% in STAD (403), 7.5% in COAD (313), and 21.4% in PAAD (33), as a gene is more easily zero in *every* specimen the fewer specimens there are. Under a weaker reading, the classes rise in order (0.0%–0.1%, 0.5%–2.3%, 19.0%–31.7%, and 100% in GDC v2; Table 4). What does not depend on the cohort size is the complete overlap class itself, which is at unity in GDC v2, even in the 1,138 BRCA specimens.

The difference between the two generations at this threshold is informative because it follows two counting rules. Of the 239 gene-by-cohort instances with complete exonic overlap in GENCODE v22, 194 have all of their exonic bases claimed by a **single** partner, and every one of them is zero in every specimen; of the 45 that need two or more partners to be covered, 40 have at least one non-zero specimen (*BOLA2*, *CIDEB*, *GNAT2*, *LRRC24*, *NME1-NME2*, *NPB*, *SERINC4*, *TMOD4*, *TNFSF12-TNFSF13*). `htseq-count -m intersection-nonempty` assigns a read to the intersection of the gene sets over the positions it covers; thus, a read that falls under *A* and *B* in one stretch and under *A* and *C* in another intersects with *A* alone and is counted; with a single covering partner, no such stretch exists. STAR `--quantMode GeneCounts` has no intersection step and discards the read whenever it touches more than one gene, which is why in GDC v2, both classes behave identically: all 246 instances with complete v36 overlap are zero everywhere, whether one partner covers the gene or several. One gene, GNAT2, is completely overlapped in v22, borderline under the HTSeq rule, and was robustly detected in GDC v2 (99%–100% of specimens) because its v36 overlap is below the threshold; the annotation, not the gene, decides.

Complete exonic overlap in the annotation against which a generation was counted is thus not a risk factor for GDC v2 but a determination of it, and for GDC v1, a determination wherever a single partner does the covering.

This also explains the features of the two-generation picture that would otherwise appear as a weak classifier. Requiring a gene to be undetectable in **both** GDC generations, while classifying it by the current annotation alone, makes only 39%–43% of genes with at least 90% v36 overlap appear lost (against 0.013%–0.098% of the rest; Fisher’s exact test, two-sided, odds ratio 733– 5,555, *P* ≤ 1.1 × 10^-70^; Table 5). The genes that escape that definition are not counterexamples: of the 74 gene-by-cohort instances in which a complete-overlap gene is not lost in both generations, each one is non-zero in GDC v1 and zero in the current generation, and 61% of them had a unique exonic sequence under GENCODE v22 and lost it only with v36. They are genes caught in the act of becoming unmeasurable, not genes that contradict the rule.

**Table 5.** Enrichment of the two-generation loss by the GDC’s nested-gene lists and by exonic overlap.

| **Cohort** | **Specimens** | **Legacy-detected genes** | **Nested, same strand** | **Nested, opposite strand** | **Not nested** | **Odds ratio (nested vs not)** | **\*P\*** | **Odds ratio (≥90% exonic vs rest)** | **\*P\*** |
|---|---|---|---|---|---|---|---|---|---|
| BRCA | 1,138 | 15,049 | 22/37 (59.5%) | 10/15 (66.7%) | 14/14,997 (0.09%) | 7,614 | 6.3 × 10<sup>-85</sup> | 1,182 | 6.2 × 10<sup>-78</sup> |
| COAD | 313 | 14,919 | 22/36 (61.1%) | 9/14 (64.3%) | 19/14,869 (0.13%) | 5,285 | 5.9 × 10<sup>-80</sup> | 733 | 1.2 × 10<sup>-72</sup> |
| LUAD | 543 | 15,081 | 21/35 (60.0%) | 9/14 (64.3%) | 14/15,032 (0.09%) | 7,169 | 4.9 × 10<sup>-80</sup> | 961 | 1.1 × 10<sup>-70</sup> |
| PAAD | 33 | 15,377 | 22/36 (61.1%) | 11/16 (68.8%) | 21/15,325 (0.14%) | 5,208 | 2.5 × 10<sup>-84</sup> | 764 | 7.5 × 10<sup>-79</sup> |
| STAD | 403 | 15,273 | 28/43 (65.1%) | 10/15 (66.7%) | 5/15,215 (0.03%) | 29,689 | 1.9 × 10<sup>-106</sup> | 5,555 | 3.7 × 10<sup>-92</sup> |

Counts are legacy-detected genes that become undetectable in both GDC generations, out of all legacy-detected genes of that class. Odds ratios and *P* values are from Fisher’s exact test (two-sided): nested on either strand versus not nested, and at least 90% exonic overlap in GENCODE v36 versus the rest.

The GDC distributes two lists of genes “completely encompassed by other genes,” which it notes “will likely display a value of zero” [41]; of those, 36–44 genes per cohort are detected in the legacy release, and 92.0%–93.5% of the same-strand list and 75.0%–78.6% of the strandless-only list are lost in both generations against 0.03%–0.13% of genes on neither list (Table 5). List membership is thus a strong predictor in its own right, and the two classifications agree closely; every one of the 40 lost genes on the lists also carries at least 90% exonic overlap in GENCODE v36. The exonic criterion is the wider of the two (among the genes lost here, it adds six further genes at that threshold and ten at 50%) for the reasons given under *Affected genes form a short, enumerable list* and *Two mechanisms, separable by the exonic criterion*.

#### Affected genes form a short, enumerable list

Because the effect follows the annotation, the affected genes can be listed. Seventy genes were undetectable in both GDC generations in at least one cohort and 32 in all five (Table 6; the full list with the per-cohort pattern is Supplementary Table S1). Of the 32, 17 are protein-coding, 10 are small nucleolar or small Cajal body-specific RNAs, four are pseudogenes, and one is a ribozyme (*RPPH1*, the RNase P RNA). All 32 share at least 99% of their exonic bases with another gene in GENCODE v36, with the lowest being *GDF1* at 99.1%. Their legacy expression is not marginal: the median legacy value of the top entries runs from hundreds to nearly two thousand normalized counts, which is within the range any analysis would treat as a solid expression. *TOMM6*, *PIGY*, *OVCA2*, *SFT2D3*, *TREX1*, *RSC1A1*, *TNFRSF6B*, *PTPRCAP*, and *ICAM4* all appear in the cancer literature, and in the GDC-era matrices, all of them are indistinguishable from genes that are genuinely silent.

**Table 6.** The 32 genes lost in all five cohorts.

| Gene symbol | Biotype | Exonic overlap | GDC nested list | Legacy median |
|---|---|---|---|---|
| TOMM6 | Protein coding | 100.0% | Same strand | 1,940 |
| PIGY | Protein coding | 100.0% | Same strand | 1,325 |
| BOLA2 | Protein coding | 100.0% | — | 1,028 |
| OVCA2 | Protein coding | 100.0% | Same strand | 493 |
| SFT2D3 | Protein coding | 100.0% | Opposite strand | 364 |
| SNORA8 | snoRNA | 100.0% | Same strand | 316 |
| TREX1 | Protein coding | 100.0% | Same strand | 314 |
| TNFRSF6B | Protein coding | 100.0% | Same strand | 281 |
| RSC1A1 | Protein coding | 100.0% | Same strand | 200 |
| SDHAP2 | Pseudogene (transcribed, unprocessed) | 100.0% | Same strand | 148 |
| PTPRCAP | Protein coding | 100.0% | Same strand | 143 |
| CEMP1 | Protein coding | 100.0% | Opposite strand | 128 |
| ANKRD36BP1 | Pseudogene (transcribed, processed) | 100.0% | Opposite strand | 108 |
| DND1 | Protein coding | 100.0% | Opposite strand | 77 |
| TMED10P1 | Pseudogene (processed) | 100.0% | Opposite strand | 52 |
| MSMP | Protein coding | 100.0% | Opposite strand | 33 |
| SNORD1C | snoRNA | 100.0% | Same strand | 24 |
| SNORD116-28 | snoRNA | 100.0% | Same strand | 21 |
| PGAM2 | Protein coding | 100.0% | Opposite strand | 19 |
| RPPH1 | Ribozyme | 100.0% | Same strand | 15 |
| ICAM4 | Protein coding | 100.0% | Opposite strand | 11 |
| NPFF | Protein coding | 100.0% | Same strand | 6.0 |
| SCARNA17 | scaRNA | 100.0% | Same strand | 5.5 |
| SNORD116-20 | snoRNA | 100.0% | Same strand | 5.1 |
| SLC7A5P2 | Pseudogene (transcribed, processed) | 100.0% | Same strand | 4.1 |
| SCARNA2 | scaRNA | 100.0% | Same strand | 2.8 |
| GDF1 | Protein coding | 99.1% | Same strand | 1.7 |
| SCARNA16 | scaRNA | 100.0% | Same strand | 0.7 |
| SNORD10 | snoRNA | 100.0% | Same strand | 0.4 |
| CPLX3 | Protein coding | 99.7% | — | 0.4 |
| SNORA67 | snoRNA | 100.0% | Same strand | 0.4 |
| SNORA57 | snoRNA | 100.0% | Same strand | 0.3 |

Ordered by legacy expression; the legacy median is the median across cohorts of the per-cohort median value (normalized\_count, tumor and normal pooled). Exonic overlap, given to one decimal place, is the fraction of the gene’s exonic bases that are also exonic in another gene in GENCODE v36; the nested-list class is the GDC’s. The full list of 70 genes, including those lost in a subset of cohorts, is Supplementary Table S1; the same analysis with a stricter legacy threshold is Supplementary Table S5.

#### Two mechanisms, separable by the exonic criterion

Of the 70 genes, 40 are on the lists the GDC publishes of “nested” genes, those completely encompassed by another gene and expected to show zero values, whereas 30 are not on these lists. The exonic criterion divides the latter into two categories. Ten of the 30 share at least half of their exonic bases with another gene (*BOLA2*, *INS-IGF2*, and *LRRC24* at 100%; *CPLX3* at 99.7%; *HAPLN4* at 97.3%; *KLK9* at 96.3%; and *APOC4*, *TSSK1B*, *RFPL1*, and *TPTE* between 72% and 83%) and are therefore the same phenomenon. They are absent from the GDC’s lists because those lists require one gene span to be completely encompassed by another, whereas exonic overlap can be complete without that being true. *BOLA2*, for instance, has all of its exonic bases claimed between four partners, chiefly the *BOLA2*–*SMG1P6* readthrough, and no single partner encompasses it. This is the first reason for preferring the exonic criterion; those 10 genes, and the 20 discussed next, are listed with their overlapping fractions in both annotations in Supplementary Table S3. Two of the twenty are classified differently by GENCODE v22, where *HBG1* reaches 71.8% and *TBC1D3H* 66.1%; the split below is therefore the v36 one, used because the two-generation definition of loss behind Tables 5 and 6 is itself applied on the v36 classification; the per-generation analysis of Fig. 2 and Table 4 classifies each generation by its own annotation.

The remaining 20 genes all fall below 50% exonic overlap in GENCODE v36 (*LIMS3* at 48.7% is the highest, followed by *CGB1* at 42.5% and *BAGE2* at 35.0%) and are a recognizable set: *TBC1D3*, *TBC1D3C*, and *TBC1D3H*; *POM121L1P*, *POM121L8P*, and *POM121L10P*; *DEFA1B*, *AMY1A*, *PGA3*, *PGA4*, *HBG1*, *CGB1*, *NXF2*, *ADAM6*, *BAGE2*, *AQP7P3*, *CXADRP2*, *CXorf49B*, *TP53TG3B*, and *LIMS3*, all members of near-identical multi-copy families. These are lost for a different reason: reads from such families map equally well to several loci, STAR GeneCounts counts only uniquely mapped reads, and the GDC v1 workflow discards alignments below MAPQ 10, whereas RSEM distributes multi-mapping reads over compatible transcripts by expectation maximization [7]. Gao et al. reported the same mechanism, finding the largest legacy-versus-harmonized discrepancies among homologous gene families [15]. Ambiguity between annotations and ambiguity between paralogs are distinct causes with the same visible consequence; the exonic criterion separates them, and a sequence-identity criterion that would quantify the second is outside the scope of this report. *BOLA2* belongs to both groups (its exons are fully claimed by annotated neighbors and are nearly identical to *BOLA2B*) and is a reminder that the two causes can coincide.

### A worked example: *TREX1*

*TREX1* encodes the major mammalian 3′→5′ DNA exonuclease. Loss-of-function mutations cause Aicardi–Goutières syndrome [43], and the enzyme restrains cell-intrinsic, nucleic-acid-driven interferon responses [44], which makes its tumor expression of obvious interest in immuno-oncology.

In the legacy release, *TREX1* was detected in 100% of the specimens from all five cohorts, with a tumor median of 281–347 normalized counts. In both GDC generations, it was zero in 100% of the specimens in all five cohorts (Fig. 3a). The transition was not gradual or cohort specific; it was complete in both re-processed generations (Fig. 3b).

**Figure 3.**
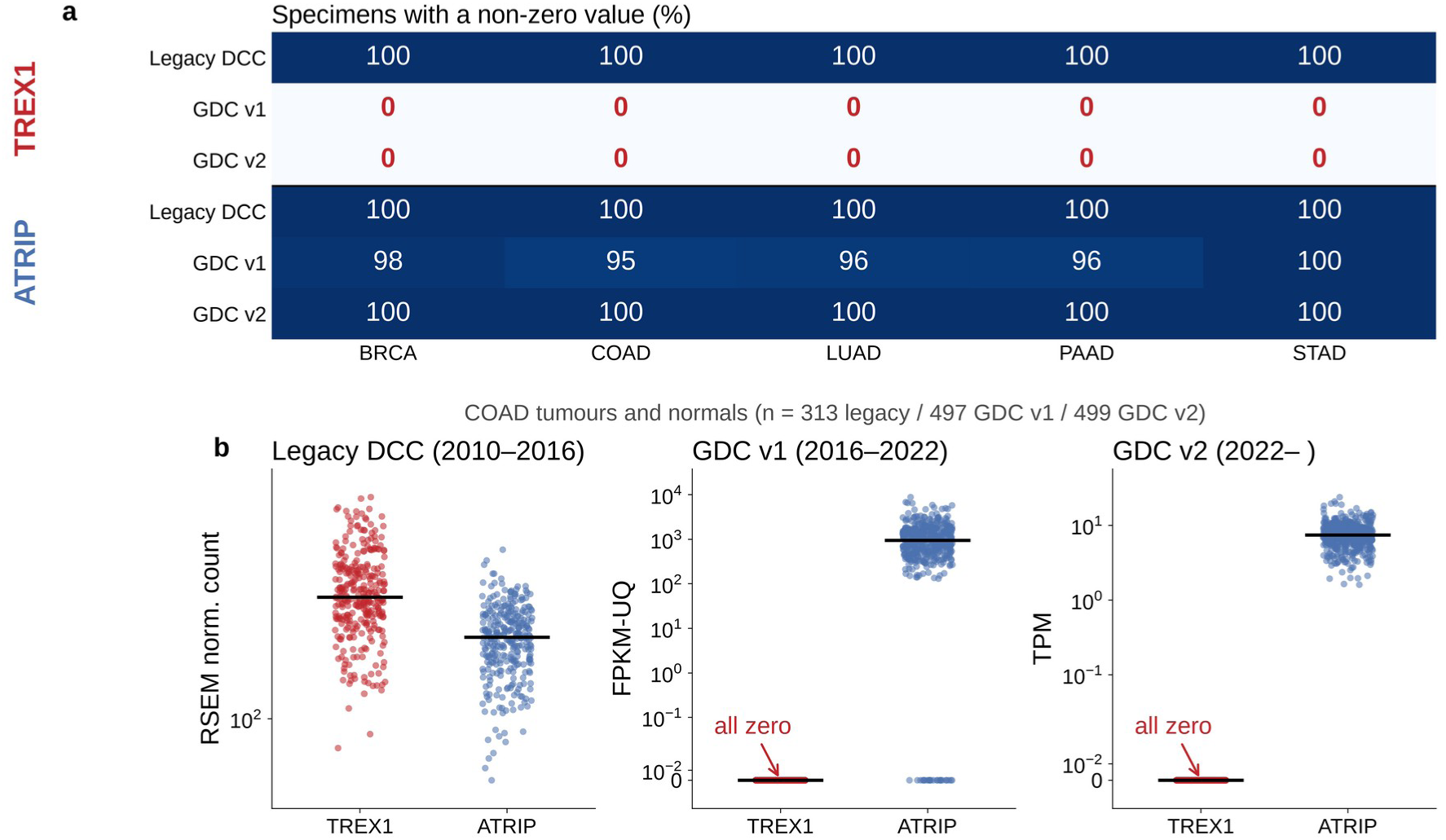
*TREX1* is lost in both GDC generations, whereas *ATRIP* is not. (**a**) Percentage of specimens with a non-zero value for *TREX1* and *ATRIP*, per generation and cohort. (**b**) Per-specimen values in COAD (tumor and normal pooled) in each generation; the *y* axis is symmetric-log, and exact zeros are visible. Units differ between generations by construction.

An obvious alternative explanation is that the locus is poorly covered after realignment to GRCh38 and is excluded by reads that remain at the locus. In GENCODE v36, *TREX1* (chr3:48,465,811–48,467,645, plus strand, 1.8 kb) lies inside the 3′ end of *ATRIP* (chr3:48,446,710–48,467,645, plus strand, 20.9 kb), and **all 1,448 of its exonic bases are also exonic in *ATRIP** *(Fig. 4), which retains 4.0 kb of exonic sequence of its own and is detected in 100% of specimens in GDC v2.* SHISA5 *lies on the opposite strand with its 3′ end 153 bp beyond the end of* TREX1 *and does not overlap in this annotation. GENCODE v22, against which GDC v1 was counted, describes the locus differently and arrives at the same place: there* ATRIP *ends 95 bp before* TREX1 *begins and does not overlap, but a third gene, the* ATRIP*–*TREX1 *readthrough ENSG00000280804 (itself labeled* TREX1*), spans both and claims all 1,835 exonic bases of* TREX1 *while leaving* ATRIP *21% of its own. The GDC v1 matrices behave accordingly: in COAD, the readthrough is non-zero in 485 of the 497 specimens of that generation and* ATRIP *in 474, whereas* TREX1 *is zero in all 497 (the comparison itself uses the 313 specimens common to all three generations). In each generation, therefore, every read falling in* TREX1* is compatible with two annotated genes, is recorded as ambiguous, and is discarded, and the gene that keeps the reads is whichever neighbor retains an exonic sequence of its own.

**Figure 4.**
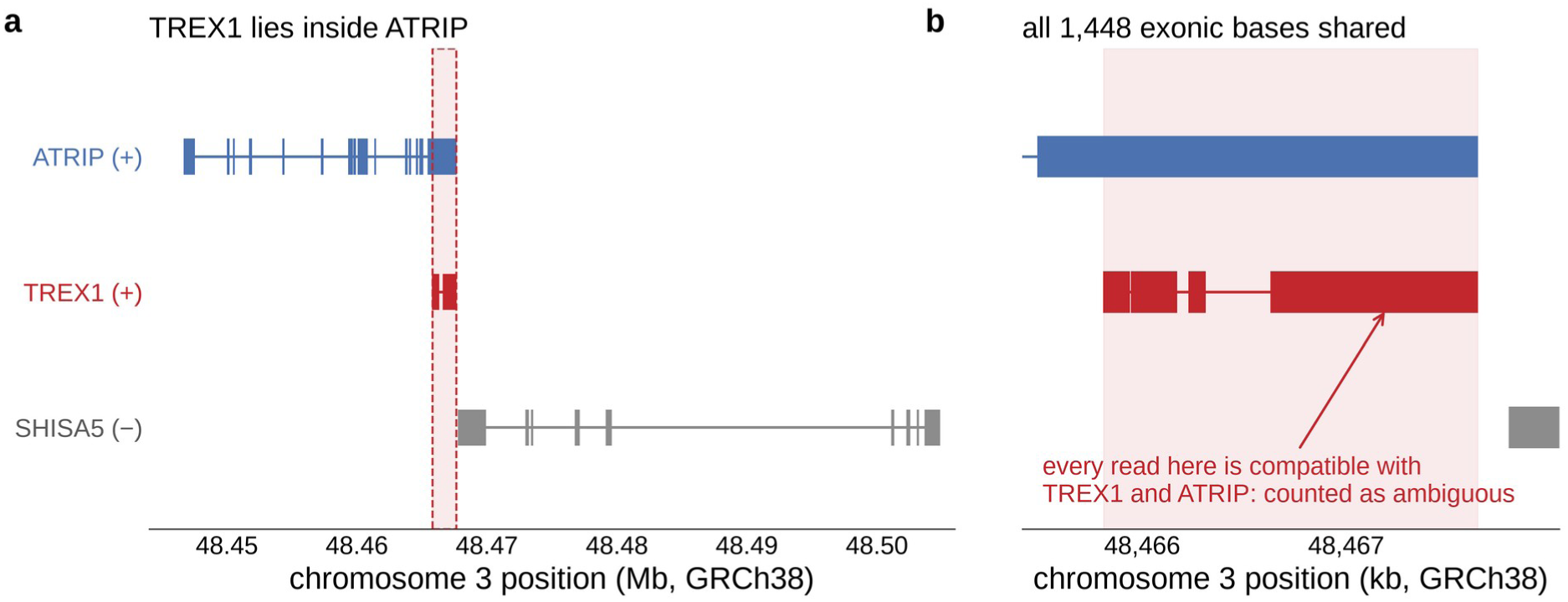
All exonic bases of *TREX1* are also exonic in *ATRIP*. Exon structure at the *TREX1* locus in GENCODE v36. For each gene, the boxes are the union of the exons of all its annotated transcripts, which is the set the counters use, lines are the intervening introns, and the shaded band is the *TREX1* gene span. (**a**) *TREX1* (1.8 kb, plus strand) lies inside the 3′ end of *ATRIP* (20.9 kb, plus strand); *SHISA5* lies on the opposite strand, beginning 153 bp downstream of TREX1, and does not overlap it in this annotation. (**b**) Enlargement of the dashed interval: all 1,448 exonic bases of *TREX1* fall inside one *ATRIP* exon; thus, every read there is compatible with two annotated genes and is discarded as ambiguous, whereas *ATRIP* keeps the reads falling in its remaining 4.0 kb of exonic sequence that no other gene claims. In GENCODE v22, the annotation of GDC v1, *ATRIP* ends 95 bp before *TREX1* begins, and the gene that claims all of *TREX1* is instead the *ATRIP*–*TREX1* readthrough ENSG00000280804 (not drawn).

RSEM, used for the legacy release, resolves the same reads differently, distributing ambiguous reads over compatible transcripts by expectation maximization [7]; therefore, the nested gene receives a non-zero estimate. The two answers are not a correct and an incorrect one; they answer different questions, and the difference becomes visible only when detection rather than magnitude is changed. It follows, and is worth stating plainly, that the legacy values at these loci should not be treated as a gold standard either; they are estimates conditional on a model of how ambiguous reads are shared.

#### The counting rule alone reproduces the loss on real reads

The mechanism was then tested directly. Two public polyA-selected RNA-seq libraries, a cell line (HepG2) and normal tissue (sigmoid colon), were each aligned once per annotation, and each alignment was counted so that the counting rule was the only thing that differed between the five GENCODE v36 conditions; a sixth condition repeated the GDC v1 rule entirely under GENCODE v22, alignment included, because the annotation of that generation describes this locus differently (*Direct test of the counting rule on public RNA-seq*; Fig. 5, Supplementary Table S4).

**Figure 5.**
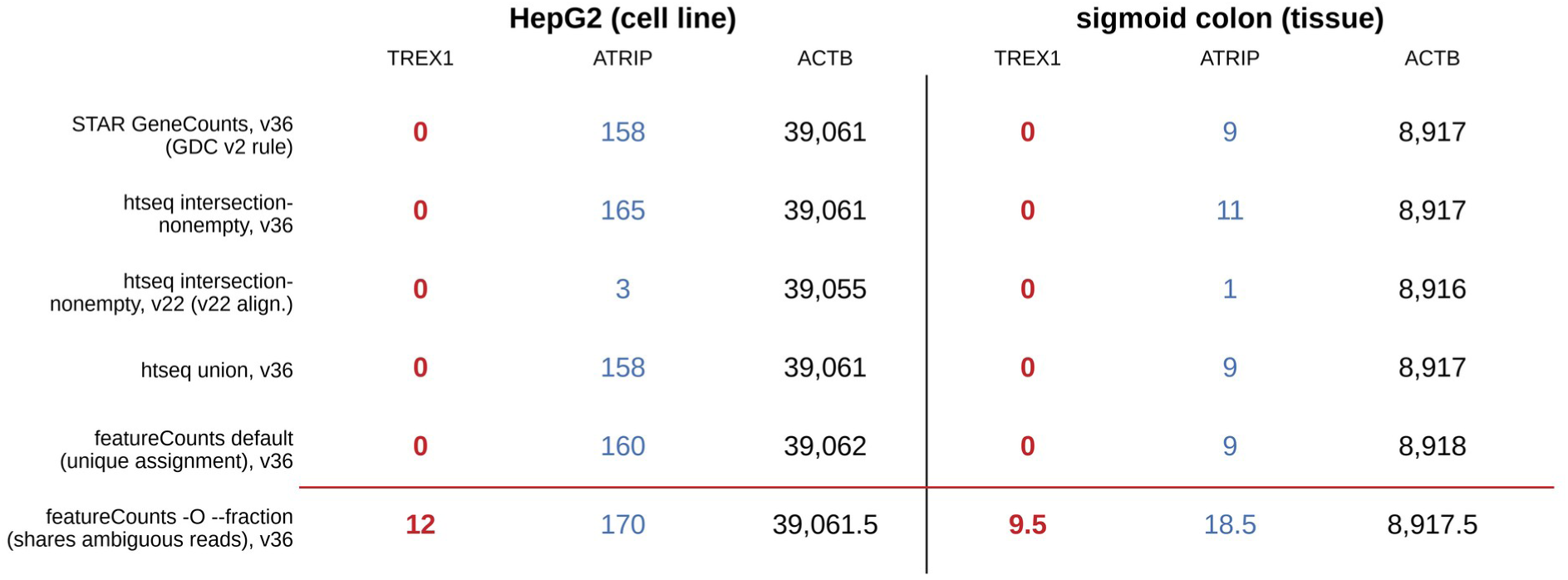
The counting rule alone turns TREX1 into a zero. Read pairs assigned to *TREX1*, *ATRIP* (the gene that contains it in GENCODE v36), and *ACTB* (a gene with no overlapping neighbor) in two public polyA-selected RNA-seq libraries, HepG2 and sigmoid colon. Each library was aligned once per annotation, and the same alignment was counted under each rule; only the counting rule, and for the third row, the annotation and its alignment, differ between rows. Rules above the red line discard reads compatible with more than one gene; the rule below it divides them between the overlapping features, which is where the fractional values come from. Under GENCODE v22, the locus carries three mutually overlapping genes, and the counts of all three collapse in these libraries (the *ATRIP*–*TREX1* readthrough receives nine pairs in HepG2 and 0 in colon); the TCGA GDC v1 matrices under the same annotation keep *ATRIP* and the readthrough non-zero and *TREX1* at zero, whereas in these two libraries (16.9 and 27.5 million read pairs), all three collapse.

*TREX1* received exactly zero read pairs under every rule that discards reads compatible with more than one gene: STAR `--quantMode GeneCounts` (the GDC v2 rule), `htseq-count -m intersection-nonempty` against GENCODE v36 (the GDC v1 rule), GENCODE v22, `htseq-count -m union`, and `featureCounts` with its default unique assignment. It received a non-zero count (12.0 pairs in HepG2 and 9.5 in the colon) only under `featureCounts -O -- fraction`, which divides an ambiguous read between the features it overlaps instead of discarding it. Across the five v36 conditions, *ATRIP* stayed in the same range (158–170 in HepG2, 9–18.5 in colon), and *ACTB*, a gene with no overlapping neighbor, was invariant to within one read pair across the five GENCODE v36 conditions (39,061–39,062 in HepG2 and 8,917–8,918 in the colon). Therefore, the counting rule, applied to identical alignments, is sufficient on its own to turn a detected gene into zero, and sharing ambiguous reads is sufficient to restore it.

Two details of this experiment are worth noting. The counters’ own tallies show where the reads go: `htseq-count` placed 718,860 pairs (HepG2) and 88,247 (colon) in the `ambiguous` bin under intersection-nonempty, and 1,532,486 and 2,041,978 under union, indicating that ambiguity is a library-wide phenomenon that is not specific to this locus. Union mode is `htseq-count`’s own default and is neither generation’s rule; it is shown because it implements the same discard- on-ambiguity behavior as STAR `--quantMode GeneCounts` and, as expected, returns per-gene counts and ambiguous totals identical to it in these libraries. Under GENCODE v22, the three mutually overlapping genes at the locus (*ATRIP*, *TREX1*, and the readthrough) all collapse in this library, to 3, 0, and 9 pairs in HepG2 and to 1, 0, and 0 in the colon when the libraries are re-aligned and counted entirely under v22 because *ATRIP*’s remaining 21% of unique exonic sequences received too few pairs in these two libraries, which carry 16.9 million (HepG2) and 27.5 million (colon) read pairs, to stay above zero. TCGA GDC v1 matrices under the same annotation leave *ATRIP* and the readthrough non-zero and *TREX1* at zero, as above, and the depth of these libraries cannot be recovered from the distributed normalized matrices. Therefore, we do not attribute the difference to depth beyond noting it. The annotation version changes which of the neighbors keeps the reads and not whether the contained gene has lost them.

#### Sensitivity to the thresholds and to the legacy values

Two conventions enter the two-generation comparison: what counts as detected in the legacy release and what counts as undetectable afterwards. Repeating the comparison under the four pairs of thresholds changed the headline percentage by a few points and nothing else (classification by at least 90% exonic overlap in GENCODE v36, lost in both GDC generations).

- non-zero in >50%/≤10% (as reported): 39.1%–42.9% of genes with at least 90% exonic overlap, 0.013%–0.098% of the rest
- non-zero in >80%/≤2%: 34.2%–37.8% of genes with at least 90% exonic overlap, 0.000%– 0.021% of the rest
- non-zero in >90%/exactly zero: 33.8%–35.6% of genes with at least 90% exonic overlap, 0.000%–0.007% of the rest
- non-zero in >30%/≤20%: 43.9%–47.5% of genes with at least 90% exonic overlap, 0.064%– 0.275% of the rest

The per-generation result in Fig. 2 is reported under the strictest reading (zero in every specimen), and only the complete-overlap class in GDC v2 is insensitive to this choice, being 100% either way. The classes below it are not: relaxing “zero everywhere” to “non-zero in at most 10% of specimens” raises the 90%–<100% class in GDC v2 from 0.0%–21.4% to 19.0%–31.7%, and the complete-overlap class in GDC v1 from 82.2%–85.4% to 86.7%–88.7%. The results are summarized in Table 4.

Because RSEM returns fractional values, a legacy value just above zero is also a weaker claim of expression than a non-zero integer count. Requiring, in addition, a legacy median above one unit leaves the conclusion unchanged (35.0%–37.3% of genes with at least 90% exonic overlap lost, against 0.007%–0.061% of the rest (Supplementary Table S5)) while removing eight of the 18 small nucleolar and small Cajal body-specific RNAs from the gene list, whose legacy medians were below one count. Therefore, the effect is not an artifact of treating the RSEM fractional values as detection, but the identity of the genes at the bottom of Table 6 depends on that convention.

## Discussion

The result reframes what a version difference in a public resource means. Re-processing a dataset is usually discussed as a change in values: different units, different normalizations, and correlations slightly below one. For genes whose exons are claimed by a neighbor, re-processing TCGA changed the membership of the measurable set. Genes that were non-zero in most or all specimens of the legacy release become undetectable in the GDC-era matrices, 32 of them in all five cohorts. Of those 32, all but three have no non-zero value in any specimen of either GDC generation; the exceptions are *CPLX3*, *GDF1*, and *BOLA2*, whose residual signal reaches 4.0%, 0.6%, and 0.2% of the specimens, respectively. Gene-level absence in one generation and robust presence in another are not subtle discrepancies that can be absorbed by averaging or rank-based methods.

Two practical consequences are as follows. First, a negative result for one of these genes in the GDC-era data is uninterpretable without checking the earlier generation; conversely, a positive result in legacy data should be read as an estimate that depends on read-sharing assumptions. Screening for silenced genes, comparisons of detection across tissues, and any pipeline that filters genes by detection rate will behave differently in the three generations; the difference is concentrated in a class of genes known before any data are touched. Second, the fix at the analysis level is available: quantification that models transcript-level assignment and then aggregates to genes [20,21] does not discard ambiguous reads, and re-processed compendia built on such methods [12,13,14] are expected to behave differently again, which is itself a reason to record the exact generation of any TCGA expression value used, by data release number rather than by the name “TCGA.”

Our findings are consistent with and extend earlier work showing that ambiguous-read handling distorts the quantification of genes in duplicated or overlapping regions [19]. The multi-copy families among our lost genes are examples of the duplicated-region case, and the exonic-overlap classifier separates them from the annotation-driven case. The findings also lie alongside rather than against the systematic legacy-versus-harmonized comparison of Gao et al. [15]. The three features of this comparison explain why the effect reported here was not visible. It was restricted to the 19,744 protein-coding genes whose identifiers could be matched unambiguously between the two releases, a selection on annotation bookkeeping rather than on read mapping; nonetheless, it removes the non-coding part of the affected class by construction, and 15 of the 32 genes lost in every cohort are non-coding. It summarizes the agreement as a per-specimen rank correlation over those 19,744 genes, a statistic that a few dozen genes going to zero cannot move. It necessarily predates the 2022 STAR-Counts re-processing; therefore, it cannot ask whether a discrepancy persists across both re-processings, which is the question that makes the present result actionable: a gene that disappears in one generation and stays absent in the next is not a transient artifact of one workflow. The same paper reports read-assignment differences of this kind from the other side: it describes them for miRNA families, where the two alignment versions distribute multi-mapping reads differently and finds the largest mRNA discordance among homologous gene families. The mechanism described here is the gene-level, annotation-driven counterpart, and it has a sharper consequence because the GDC counters discard ambiguous reads rather than redistribute them.

The behavior itself is not unknown to the data provider: the GDC’s own mRNA pipeline documentation distributes the two lists of genes “completely encompassed by other genes” and states that they “will likely display a value of zero” [41]. What has been added here are the quantitative and comparative aspects of this statement. The expectation is exact once the question is asked of one generation at a time and the gene is classified by exonic bases rather than by gene span: the 46–55 genes per cohort whose exons are entirely claimed by others in GENCODE v36 have no non-zero value anywhere in GDC v2, without exception, in five cohorts spanning a 34-fold range of specimen number. The two GDC generations differ at that threshold in a way that follows from their counting rules rather than from the annotation alone: the loss is reproduced on public reads by changing nothing but the counting rule, and the check is now available in a browser rather than requiring re-analysis.

### Limitations

The tool currently covers five cohorts, and the comparison presented here is therefore not pan-cancer. Extending it to all TCGA cohorts is in progress. The counting experiment in Fig. 5 was run on two public libraries rather than on TCGA alignments, which are controlled access and would have to be re-counted at scale, establishing that the counting rule is sufficient to produce the loss in reads of the same kind, and not that every TCGA zero arose that way. The exonic-overlap fractions are computed from the annotation each generation was counted against; therefore, the classification is annotation-specific by construction: a gene can be unmeasurable in one generation and measurable in another because the annotation changed around it, which is what the 90%–<100% class and the complete-overlap genes detected only in GDC v1 show. The detection thresholds (non-zero in more than 50% of specimens for “detected”; non-zero in at most 10% for “undetectable”) are conventions chosen for clarity; the sensitivity analysis above shows the effect does not depend on them, whereas the exact gene counts in Tables 5 and 6 do. Both overlap measures treat a gene as the union of the exons of all its annotated transcripts, as counters do; therefore, a gene whose overlap is confined to a minor isoform is classified with the rest. The legacy matrices are indexed by gene symbol; thus, cross-generation comparison required symbol-to-Ensembl mapping and genes with ambiguous symbols were dropped. The *ATRIP*–*TREX1* readthrough, which carries the symbol *TREX1* in GENCODE v22, is an example of why that mapping has to be done on identifiers. The GDC v2 matrices were downloaded at different GDC releases per cohort (Data Release 36.0 to 46.0; Table 2), which the pipeline itself does not change but which means that the specimen sets of that generation were frozen at different dates. Finally, detection rather than magnitude was compared deliberately: unit and scaling conventions differ between generations, and one cohort’s middle-generation source is distributed on a different scale from the others; therefore, magnitude comparisons across generations would confound pipeline effects with unit conventions.

### A gene with no unique exonic sequence reports which generation a resource serves

The genes described herein are both diagnostic and problematic. A resource that serves the legacy generation returns a distribution for *TREX1* in every cohort we examined, whereas a resource that serves either the GDC generation returns zero or no entry at all (Fig. 3a). Therefore, querying one such gene identifies the generation behind a resource whose interface does not state this. We queried *TREX1* in OncoDB and in the TNM plot and obtained no expression for it in either, which is what the generation each serves (Table 3) predicts; the same query in a legacy-derived resource returns the distribution that the legacy matrices contain. The genes listed in Supplementary Table S1 are usable in this manner, and the check requires one query.

This argument runs in another manner. A reader who finds a gene from that list reported as expressed in one resource and absent in another does not need to suspect either resource; the two serve different generations of the same experiment. This is also why we report the generation of every resource in Table 3 from its own data provenance statement rather than from its interface, which, in most cases, does not state this.

### Potential implications

A gene-level expression value from a re-processed public resource carries a version. For a minority of genes, the version determines whether the gene is measurable at all. We suggest that resources of this kind ship, and that tools of this kind display, an explicit per-gene flag for annotation-driven ambiguity. The list is short, computable from the annotation alone, and stable until the annotation changes. TCGA Expression Explorer implements the display side of that suggestion, and the same design applies unchanged to any other resource that has been re-processed against successive annotations.

### Planned future development

Three extensions are planned: coverage of all TCGA cohorts rather than the present five; a per-gene warning flag in the interface, computed from the annotation, which marks genes whose locus overlaps another gene before the user interprets a zero; and read-level confirmation of the mechanism on long-read data for a subset of the affected loci. Transcript-level values and programmatic endpoints for batch queries are under consideration.

## Methods

### Cross-generation comparison

The comparison used the deposited source matrices rather than the tool’s JSON documents and reproduced the specimen selection of the tool exactly. Accordingly, the numbers reported here and the distributions a user sees come from the same specimens. Columns were reduced to 15-character TCGA barcodes; codes 02, 04, 06, 07, 08, and 40 (recurrent and metastatic) were excluded first; of the remainder, codes 01, 03, 05, and 09 were taken as tumor, and any code of 10 or above was considered normal. In these five cohorts, the surviving specimens carried only codes 01 (primary solid tumor) and 11 (solid tissue normal); the remaining tumor codes were retained in the rule because the tool applies it to cohorts that contain them. Duplicate vials were handled as follows: duplicate vials of the same patient and sample type were collapsed into the first vial, and the per-cohort exclusion lists used to build the tool were applied: formalin-fixed specimens identified from the GDC sample sheet (13 in BRCA, 12 in LUAD), 19 specimens of non-pancreatic ductal histology, and nine pseudonormal specimens flagged by the cohort’s marker paper [29]. The analysis was restricted to specimens present in all three generations, leaving 1,138 (BRCA), 313 (COAD), 543 (LUAD), 33 (PAAD), and 403 (STAD) specimens.

Legacy rows are labeled by gene symbols; GDC v2 rows carry both Ensembl gene ID and symbols, and GDC v1 rows carry Ensembl gene IDs only. Symbols were therefore mapped to Ensembl gene IDs using the symbol-to-identifier correspondence of the GDC v2 matrix, keeping only symbols that map to exactly one identifier, where a matrix still carried the same label on more than one row, and the first occurrence was kept. This yielded 17,352 genes that were comparable across all three generations in every cohort.

For each gene and generation, we computed the fraction of compared specimens with a value greater than zero. A gene was called *detected* in the legacy generation when that fraction exceeded 0.5. Two outcomes that are not interchangeable have been reported. The per-generation outcome in Fig. 2 and Table 4 is *zero in every compared specimen* of that generation, the strictest available reading; the two-generation outcome used for Tables 5 and 6 and for the supplementary gene lists is *lost*, meaning detected in the legacy generation and non-zero in at most 10% of the specimens in both GDC generations. Table 4 lists the per-generation counts for both the readings.

These genes are classified into two categories. The primary classifier is the fraction of a gene’s exonic bases that are also exonic in at least one other gene. For each gene, the exons of all its transcripts were merged into a set of intervals; the same was done for every other gene, and the classifier is the number of the gene’s exonic bases covered by the union of all other genes’ merged exons divided by its own exonic length. The union is taken irrespective of the strand because the GDC counts all reads as unstranded [41] and over all partners jointly rather than one partner at a time, because STAR’s GeneCounts mode discards a read as soon as it is compatible with a second gene, whatever the gene is. The same union is used for GDC v1 for comparability, but it is not equivalent to the counter’s rule: `htseq-count -m intersection-nonempty` assigns a read to the intersection of the gene sets along its length and can therefore still count a read that touches a second gene over part of its span, which is why the two generations differ at complete overlap. It was computed twice, from the GENCODE v36 comprehensive annotation on the reference chromosomes [31] for GDC v2 and from GENCODE v22 for GDC v1. Each generation was classified by its own annotation; genes were grouped as below 50%, 50%–90%, 90% to below 100%, and 100%, the last being genes with no exonic bases of their own. The per-strand fractions were retained for reference and are reported in the deposited table. The second classifier uses the two lists the GDC distributes with its mRNA analysis pipeline documentation, `overlap.gene.stranded.tsv` and `overlap.gene.strandless.tsv` [41], which enumerate genes completely encompassed by another gene and for which the GDC states will likely show a value of zero. The stranded list (962 genes) is a subset of the strandless list (1,726 genes), and we refer to members of the stranded list as same-strand nested and to the 764 genes present only in the strandless list as opposite-strand nested. Those lists carry GENCODE v22-era entries; therefore, they describe the annotation of GDC v1. Membership was transferred to our gene table on the versionless Ensembl gene identifier rather than on the symbol, because the lists carry GENCODE v22-era names (*RPPH1*, for example, appears as `RNaseP_nuc` under the same identifier), and symbols are neither stable across releases nor unique.

Enrichment of loss was tested per cohort using Fisher’s exact test (two-sided), comparing genes with at least 90% exonic overlap against the rest of the legacy-detected genes and, for the second classifier, nested against non-nested genes. Two sensitivity analyses were run, and the whole comparison was repeated under four pairs of detection thresholds, requiring a legacy median above one `normalized_count` unit in addition to the detection rule, because RSEM assigns fractional values, and a legacy value just above zero is a weaker claim of expression than a non-zero count.

### Direct test of the counting rule on public RNA-seq

Because the aligned TCGA sequence data have controlled access, and re-counting them at scale is impractical, the counting step itself was tested on public data. Two polyA-selected, paired-end RNA-seq experiments from ENCODE were used, one cell line (HepG2, ENCSR985KAT) and one normal tissue (sigmoid colon, ENCSR999ZCI); for each, one replicate’s read pairs were aligned with STAR 2.7.11b to GRCh38 with the GENCODE v36 annotation supplied at mapping, and that alignment was counted in five ways with HTSeq 2.0.5 and Subread 2.0.6; hence, only the counting rule differs between those five conditions: STAR `--quantMode GeneCounts` (the GDC v2 rule); `htseq-count -m intersection-nonempty -s no -a 10 -t exon` (the GDC v1 rule); `htseq-count -m union`; `featureCounts`, with its default unique assignment; and `featureCounts -O --fraction`, which divides a read between the features it overlaps instead of discarding it. The sixth condition repeats the GDC v1 rule entirely under GENCODE v22 (the same read pairs re-aligned with the v22 annotation supplied at mapping and counted against v22) because the annotation describes the *TREX1* locus differently. The two alignments differ negligibly in the mapping rate (86.06% against 86.07% unique in HepG2, 77.02% in colon both times). *ATRIP* served as the internal control and *ACTB* as a gene with no overlapping neighbors, and the counters’ own ambiguous read tallies were recorded.

The locus diagram (Fig. 4) was drawn from the GENCODE v36 annotation, showing the union of the exons of all its transcripts for each gene, which is what the counters see; the per-specimen values in Figs. 3a and 3b are those served by the tool itself. Figures were produced using Matplotlib. The comparison script, exonic-overlap script, and reduced test case with its expected output are in the code repository.

### Identification of comparable resources

Existing resources were identified by searching OpenAlex (searched September 18, 2026) with eight queries combining the terms *web server*, *web tool*, *portal*, *browser*, *interactive*, *visualization*, *TCGA*, *gene expression*, *RNA-seq*, *quantification pipeline*, *re-processing*, *harmonized*, *annotation version*, and *ambiguous reads*, restricted to 2010 onwards, followed by targeted searches for named resources and screening the reference lists of the hits. Records were screened on the title and abstract for resources that display TCGA gene-level expression for a user-chosen gene, or to compare TCGA processing generations. The 243 unique records screened yielded the resources listed in Supplementary Table S2. No resources displaying more than one quantification generation per gene were found.

### Software used

The comparisons and figures were produced using Python 3.11, pandas [45], NumPy [46], SciPy [47], and Matplotlib [48]. The exact versions are pinned in `requirements.txt` in the source repository [25]. The preprocessing pipeline also uses mygene [49] for identifier resolution. The interface renders with Plotly.js 2.35.2 [50]. The counting experiments used STAR 2.7.11b [8], HTSeq 2.0.5 [9], and Subread 2.0.6 [42], installed from bioconda. Overlap classes and locus coordinates were computed from GENCODE release 36 and release 22 annotation files [51], and the gene annotations of the three generations were GENCODE v36 (GDC v2), GENCODE v22 (GDC v1), and the UCSC hg19 gene model (legacy). The analysis corresponds to version 1.0 of the deposited data package [24] and to the repository release tagged at the time of submission.

## Availability of supporting source code and requirements

• **Project name:** TCGA Expression Explorer

• Project home page: https://5spgm.github.io/tcga-expression-explorer/ [30]; source code https://github.com/5spgm/tcga-expression-explorer [25], with the version described here archived at Zenodo, doi:10.5281/zenodo.24503264

• **Operating system(s):** Platform independent (any current web browser); preprocessing tested on Linux

• **Programming language:** JavaScript (interface), Python 3.11 (preprocessing and analysis)

• **Other requirements:** Plotly.js 2.35.2 [50]; Python ≥ 3.9 with pandas, NumPy, SciPy, Matplotlib, openpyxl, xlrd, and mygene, as pinned in requirements.txt

• **License:** MIT for the source code, as stated in the LICENSE file of the repository [25]; Creative Commons Attribution 4.0, International (CC BY 4.0) for the deposited data [24].

The underlying TCGA data are open-access and were obtained under the NCI Genomic Data Commons data use policies.

• **RRID: SCR_029343 (submitted to the SciCrunch Registry; curation pending)**

• **bio.tools ID: biotools:tcga_expression_explorer (**https://bio.tools/tcga_expression_explorer)

## Data Availability

The datasets supporting the results of this article are available in the Zenodo repository, doi:10.5281/zenodo.22670234 [24]: the source matrices for all five cohorts and three generations, specimen annotation, FFPE and flagged-specimen exclusion lists, provenance records, a machine-readable inventory, and SHA-256 checksums, released under CC BY 4.0. Five supplementary tables accompany this article: the full list of genes lost in the GDC era with their per-cohort pattern, biotype, exonic overlap, and legacy expression level (S1, also in the deposit); the resources screened for functional comparison (S2); the lost genes that the GDC’s nested-gene lists do not contain, with their exonic overlap and suggested mechanism (S3); the counts obtained from the two public libraries under each counting rule (S4); and the same loss analysis under the stricter legacy threshold (S5). The public libraries used were the ENCODE experiments ENCSR985KAT and ENCSR999ZCI. The primary data sources were the TCGA DCC Data Matrix distribution, NCI Genomic Data Commons Data Portal [23], and UCSC Xena GDC hub [10]. A snapshot of the source code in the version used here is cited as a reference [25], and upon acceptance, we will additionally archive that snapshot with a DOI in GigaDB or Software Heritage and cite it here.

## List of abbreviations

DCC: Data Coordinating Center
DR: data release
FFPE: formalin-fixed, paraffin-embedded
FPKM: fragments per kilobase of transcript per million mapped reads
FPKM-UQ: upper-quartile-normalized FPKM
GDC: Genomic Data Commons
JSON: JavaScript Object Notation
RSEM: RNA-Seq by Expectation Maximization
TCGA: The Cancer Genome Atlas
TPM: transcripts per million

## Declarations

### Ethics approval and consent to participate

Not applicable. This study used only publicly available, de-identified data from TCGA and imposed no new requirements on the human participants.

### Consent for publication

Not applicable.

### Competing interests

The authors declare that they have no competing interests.

## Funding

This work was supported by the Japan Society for the Promotion of Science (JSPS) KAKENHI, Grant-in-Aid for Scientific Research (C), grant number JP23K09154 (Principal Investigator S.T.). The funding body had no role in the design of the study; in the collection, analysis, and interpretation of data; or in writing the manuscript.

## Authors’ contributions

S.T.: conceptualization, software, data curation, formal analysis, visualization, and writing (original draft. M.I.: conceptualization, validation, supervision, writing) review, and editing. Both authors read and approved the final version of the manuscript.

## Use of AI-assisted technology

Generative AI assistance (Claude, Anthropic) [52] was used to write the plotting and analysis code and to draft parts of the manuscript text; the figures themselves were rendered by that code and were not generated as images. No data processing, computation, or statistical analysis was delegated to it. All values reported here were computed by the authors from the deposited data with the scripts in the source repository, and the authors are responsible for the content of the article.

## Acknowledgements

We thank Azusa Kubota (Kanazawa University), Seiichiro Ishihara (Hokkaido University), and Yue Zhou (University of Toyama) for testing the tool and providing suggestions. We acknowledge the TCGA Research Network, Genomic Data Commons, and the patients and families who contributed specimens.

## Notes

### Competing Interest Statement

The authors have declared no competing interest.

## References

1. Weinstein JN, Collisson EA, Mills GB, Shaw KRM, Ozenberger BA, Ellrott K, et al. The Cancer Genome Atlas Pan-Cancer analysis project. Nat Genet. 2013;45:1113–20. doi:10.1038/ng.2764.

2. Tange S, Hirano T, Idogawa M, Hirata E, Imoto I, Tokino T. MYEOV overexpression induced by demethylation of its promoter contributes to pancreatic cancer progression via activation of the folate cycle/c-Myc/mTORC1 pathway. BMC Cancer. 2023;23:85. doi:10.1186/s12885-022-10433-6.

3. Kamikokura M, Tange S, Nakase H, Tokino T, Idogawa M. Long Noncoding RNA RP11-278A23.1, a Potential Modulator of p53 Tumor Suppression, Contributes to Colorectal Cancer Progression. Cancers. 2024;16:882. doi:10.3390/cancers16050882.

4. Idogawa M, Koizumi M, Hirano T, Tange S, Nakase H, Tokino T. Dead or alive? Pitfall of survival analysis with TCGA datasets. Cancer Biol Ther. 2021;22:527–8. doi:10.1080/15384047.2021.1979845.

5. Grossman RL, Heath AP, Ferretti V, Varmus HE, Lowy DR, Kibbe WA, et al. Toward a Shared Vision for Cancer Genomic Data. N Engl J Med. 2016;375:1109–12. doi:10.1056/NEJMp1607591.

6. Wang K, Singh D, Zeng Z, Coleman SJ, Huang Y, Savich GL, et al. MapSplice: Accurate mapping of RNA-seq reads for splice junction discovery. Nucleic Acids Res. 2010;38:e178. doi:10.1093/nar/gkq622.

7. Li B, Dewey CN. RSEM: accurate transcript quantification from RNA-Seq data with or without a reference genome. BMC Bioinformatics. 2011;12:323. doi:10.1186/1471-2105-12-323.

8. Dobin A, Davis CA, Schlesinger F, Drenkow J, Zaleski C, Jha S, et al. STAR: ultrafast universal RNA-seq aligner. Bioinformatics. 2013;29:15–21. doi:10.1093/bioinformatics/bts635.

9. Anders S, Pyl PT, Huber W. HTSeq—a Python framework to work with high-throughput sequencing data. Bioinformatics. 2015;31:166–9. doi:10.1093/bioinformatics/btu638.

10. Goldman MJ, Craft B, Hastie M, Repečka K, McDade F, Kamath A, et al. Visualizing and interpreting cancer genomics data via the Xena platform. Nat Biotechnol. 2020;38:675–8. doi:10.1038/s41587-020-0546-8.

11. Colaprico A, Silva TC, Olsen C, Garofano L, Cava C, Garolini D, et al. TCGAbiolinks: an R/Bioconductor package for integrative analysis of TCGA data. Nucleic Acids Res. 2016;44:e71. doi:10.1093/nar/gkv1507.

12. Tatlow P, Piccolo SR. A cloud-based workflow to quantify transcript-expression levels in public cancer compendia. Sci Rep. 2016;6:39259. doi:10.1038/srep39259.

13. Vivian J, Rao AA, Nothaft FA, Ketchum C, Armstrong J, Novak A, et al. Toil enables reproducible, open source, big biomedical data analyses. Nat Biotechnol. 2017;35:314–6. doi:10.1038/nbt.3772.

14. Wilks C, Zheng SC, Chen FY, Charles R, Solomon B, Ling JP, et al. recount3: summaries and queries for large-scale RNA-seq expression and splicing. Genome Biol. 2021;22:323. doi:10.1186/s13059-021-02533-6.

15. Gao GF, Parker JS, Reynolds SM, Silva TC, Wang LB, Zhou W, et al. Before and After: Comparison of Legacy and Harmonized TCGA Genomic Data Commons’ Data. Cell Syst. 2019;9:24–34.e10. doi:10.1016/j.cels.2019.06.006.

16. Zhang Z, Hernandez K, Savage J, Li S, Miller D, Agrawal S, et al. Uniform genomic data analysis in the NCI Genomic Data Commons. Nat Commun. 2021;12:1226. doi:10.1038/s41467-021-21254-9.

17. Zhao S, Zhang B. A comprehensive evaluation of ensembl, RefSeq, and UCSC annotations in the context of RNA-seq read mapping and gene quantification. BMC Genomics. 2015;16:97. doi:10.1186/s12864-015-1308-8.

18. SEQC/MAQC-III Consortium. A comprehensive assessment of RNA-seq accuracy, reproducibility and information content by the Sequencing Quality Control Consortium. Nat Biotechnol. 2014;32:903–14. doi:10.1038/nbt.2957.

19. Robert C, Watson M. Errors in RNA-Seq quantification affect genes of relevance to human disease. Genome Biol. 2015;16:177. doi:10.1186/s13059-015-0734-x.

20. Soneson C, Love MI, Robinson MD. Differential analyses for RNA-seq: transcript-level estimates improve gene-level inferences. F1000Res. 2015;4:1521. doi:10.12688/f1000research.7563.1.

21. Patro R, Duggal G, Love MI, Irizarry RA, Kingsford C. Salmon provides fast and bias-aware quantification of transcript expression. Nat Methods. 2017;14:417–9. doi:10.1038/nmeth.4197.

22. GDC Data Release Notes. https://docs.gdc.cancer.gov/Data/Release_Notes/Data_Release_Notes/. Accessed 2 Oct 2026.

23. National Cancer Institute Genomic Data Commons Data Portal. https://portal.gdc.cancer.gov/. Accessed 18 Sep 2026.

24. Tange S, Idogawa M. TCGA Expression Explorer — source data (Version 1.0) [Data set]. Zenodo. 2026. doi:10.5281/zenodo.22670234.

25. Tange S, Idogawa M. TCGA Expression Explorer (Version v1.0.0) [Source code]. GitHub. 2026. https://github.com/5spgm/tcga-expression-explorer. Archived at Zenodo, doi:10.5281/zenodo.24503264.

26. Hoadley KA, Yau C, Hinoue T, Wolf DM, Lazar AJ, Drill E, et al. Cell-of-Origin Patterns Dominate the Molecular Classification of 10,000 Tumors from 33 Types of Cancer. Cell. 2018;173:291–304.e6. doi:10.1016/j.cell.2018.03.022.

27. Liu Y, Sethi NS, Hinoue T, Schneider BG, Cherniack AD, Sanchez-Vega F, et al. Comparative Molecular Analysis of Gastrointestinal Adenocarcinomas. Cancer Cell. 2018;33:721–735.e8. doi:10.1016/j.ccell.2018.03.010.

28. Thorsson V, Gibbs DL, Brown SD, Wolf D, Bortone DS, Ou Yang TH, et al. The Immune Landscape of Cancer. Immunity. 2018;48:812–830.e14. doi:10.1016/j.immuni.2018.03.023.

29. Raphael BJ, Hruban RH, Aguirre AJ, Moffitt RA, Yeh JJ, Stewart C, et al. Integrated Genomic Characterization of Pancreatic Ductal Adenocarcinoma. Cancer Cell. 2017;32:185–203.e13. doi:10.1016/j.ccell.2017.07.007.

30. TCGA Expression Explorer. https://5spgm.github.io/tcga-expression-explorer/. Accessed 18 Sep 2026.

31. Frankish A, Diekhans M, Jungreis I, Lagarde J, Loveland JE, Mudge JM, et al. GENCODE 2021. Nucleic Acids Res. 2021;49:D916–D923. doi:10.1093/nar/gkaa1087.

32. Chandrashekar DS, Karthikeyan SK, Korla PK, Patel H, Shovon AR, Athar M, et al. UALCAN: An update to the integrated cancer data analysis platform. Neoplasia. 2022;25:18–27. doi:10.1016/j.neo.2022.01.001.

33. Bartha Á, Győrffy B. TNMplot.com: A Web Tool for the Comparison of Gene Expression in Normal, Tumor and Metastatic Tissues. Int J Mol Sci. 2021;22:2622. doi:10.3390/ijms22052622.

34. Tang Z, Li C, Kang B, Gao G, Li C, Zhang Z. GEPIA: a web server for cancer and normal gene expression profiling and interactive analyses. Nucleic Acids Res. 2017;45:W98–W102. doi:10.1093/nar/gkx247.

35. Tang Z, Kang B, Li C, Chen T, Zhang Z. GEPIA2: an enhanced web server for large-scale expression profiling and interactive analysis. Nucleic Acids Res. 2019;47:W556–W560. doi:10.1093/nar/gkz430.

36. Kang YJ, Pan L, Liu Y, Rong Z, Liu J, Liu F. GEPIA3: Enhanced drug sensitivity and interaction network analysis for cancer research. Nucleic Acids Res. 2025;53:W283–W290. doi:10.1093/nar/gkaf423.

37. Tang G, Cho M, Wang X. OncoDB: an interactive online database for analysis of gene expression and viral infection in cancer. Nucleic Acids Res. 2022;50:D1334–D1339. doi:10.1093/nar/gkab970.

38. Cerami E, Gao J, Dogrusoz U, Gross BE, Sumer SO, Aksoy BA, et al. The cBio Cancer Genomics Portal: An Open Platform for Exploring Multidimensional Cancer Genomics Data. Cancer Discov. 2012;2:401–4. doi:10.1158/2159-8290.CD-12-0095.

39. Kus ME, Sahin C, Kilic E, Askin A, Ozgur MM, Karahanogullari G, et al. TCGEx: a powerful visual interface for exploring and analyzing cancer gene expression data. EMBO Rep. 2025;26:14. doi:10.1038/s44319-025-00407-7.

40. Park SJ, Yoon BH, Kim SK, Kim SY. GENT2: an updated gene expression database for normal and tumor tissues. BMC Med Genomics. 2019;12:101. doi:10.1186/s12920-019-0514-7.

41. GDC mRNA Analysis Pipeline, including the Overlapped Genes (stranded) and Overlapped Genes (unstranded) lists. https://docs.gdc.cancer.gov/Data/Bioinformatics_Pipelines/Expression_mRNA_Pipeline/. Accessed 21 Oct 2026.

42. Liao Y, Smyth GK, Shi W. featureCounts: an efficient general purpose program for assigning sequence reads to genomic features. Bioinformatics. 2014;30:923–30. doi:10.1093/bioinformatics/btt656.

43. Crow YJ, Hayward BE, Parmar R, Robins P, Leitch A, Ali M, et al. Mutations in the gene encoding the 3′-5′ DNA exonuclease TREX1 cause Aicardi-Goutières syndrome at the AGS1 locus. Nat Genet. 2006;38:917–20. doi:10.1038/ng1845.

44. Stetson DB, Ko JS, Heidmann T, Medzhitov R. Trex1 Prevents Cell-Intrinsic Initiation of Autoimmunity. Cell. 2008;134:587–98. doi:10.1016/j.cell.2008.06.032.

45. The pandas development team. pandas (Version as pinned in requirements.txt) [Software]. GitHub. https://github.com/pandas-dev/pandas. Accessed 18 Sep 2026.

46. NumPy Developers. NumPy (Version as pinned in requirements.txt) [Software]. GitHub. https://github.com/numpy/numpy. Accessed 18 Sep 2026.

47. SciPy Developers. SciPy (Version as pinned in requirements.txt) [Software]. GitHub. https://github.com/scipy/scipy. Accessed 18 Sep 2026.

48. Matplotlib Development Team. Matplotlib (Version as pinned in requirements.txt) [Software]. GitHub. https://github.com/matplotlib/matplotlib. Accessed 18 Sep 2026.

49. BioThings Team. mygene.py (Version [version]) [Software]. GitHub. https://github.com/biothings/mygene.py. Accessed 18 Sep 2026.

50. Plotly Technologies Inc. plotly.js (Version 2.35.2) [Software]. GitHub. https://github.com/plotly/plotly.js/releases/tag/v2.35.2. Accessed 18 Sep 2026.

51. GENCODE Release 36 (GRCh38.p13) and Release 22 (GRCh38) comprehensive gene annotation. https://www.gencodegenes.org/human/release_36.html and https://www.gencodegenes.org/human/release_22.html. Accessed 18 Sep 2026.

52. Anthropic. Claude [large language model]. Responses to queries made by S. Tange, August– October 2026. Claude.ai. https://claude.ai/.

